# Simultaneous assessment of human genome and methylome data in a single experiment using limited deamination of methylated cytosine

**DOI:** 10.1101/2023.06.16.545253

**Authors:** Bo Yan, Duan Wang, Laurence Ettwiller

## Abstract

Multi-omics requires concerted recording of independent information, ideally from a single experiment. In this study, we introduce RIMS-seq2, a high-throughput technique to simultaneously sequence genomes and overlay methylation information while requiring only a small modification of the experimental protocol for high throughput DNA sequencing to include a controlled deamination step. Importantly, the rate of deamination of 5mC is negligible and thus, do not interfere with standard DNA sequencing and data processing. Thus, RIMS-seq2 libraries from whole or targeted genome sequencing show the same germline variation calling accuracy and sensitivity as compared to standard DNA-seq. Additionally, regional methylation levels provide an accurate map of the human methylome.

## Introduction

Cytosine methylation is the main epigenetic DNA modification found in higher Eukaryotes. In humans it occurs mainly in CpG dinucleotide context with the help of DNA methyltransferases (DNMTs) which transfer a methyl group to a cytosine residue to form 5-methylcytosine (5mC). Methylation of cytosine is involved in various biological processes, including regulation of gene expression and chromatin structure. Additionally, abnormal methylation patterns have been found to play a significant role in disease progression and carcinogenesis (Jones and Baylin 2007). Accordingly, methylation of cytosine can be used as universal biomarkers for the diagnosis of disease, responses to therapeutic interventions and prognosis (Hulbert et al. 2017) demonstrating its usage for noninvasive detection of conditions. Furthermore DNA methylation is a fairly accessible biomarker due to its low sensitivity to experimental handling.

Techniques for the identification of DNA methylation can be grouped based on the properties used to discriminate between methylated and unmethylated cytosines, namely enzymatic digestion, affinity enrichment, and enzymatic or chemical conversion. The most commonly used techniques rely on the conversion of cytosine to uracil followed by either hybridization of the converted sequence to methylation arrays or sequencing of the whole-genome (WGBS) or of a reduced-representation (RRBS). Converting all cytosines severely reduces sequence complexity and therefore these conversion-based techniques have a single aim, namely to identify methylation. Recently, TET-assisted pyridine borane sequencing (TAPS) (Liu et al. 2019) allows the conversion of only methylated cytosine which significantly improves mapping and coverage.

Recently, we and others have used the redundancy of the double stranded DNA to identify converted cytosine and thus, obtain DNA methylation information and genomic variants simultaneously (Yan et al. 2022)(Liang et al. 2021)(Füllgrabe et al. 2023). While this setup allows for dual readouts in a single dataset, the experimental application of these techniques proved to be a significant departure from standard library preparation as the procedure involves linking double stranded DNA together.

Chemical or fortuitous deamination can also be used as means to identify cytosine methylation. For example, Gokhman et al. elegantly harness DNA damage resulting from the natural degradation processes of inappropriately stored DNA to identify methylated and unmethylated cytosines in ancient DNA (Gokhman et al. 2014). Similarly, we developed RIMS-seq, a new method to identify methylase specificity in bacteria (Baum et al. 2021). To perform RIMS-seq, genomic DNA is subjected to a limited heat-alkaline treatment step that induces a deamination of a fraction of m5C. While unmethylated cytosines are also deaminated, they are effectively eliminated during the amplification step due to the usage of a proofreading polymerase that stalls at dU sites. Thus, only the deaminated m5C results in C to T transition in sequencing reads. m5C sites are therefore identified by virtue of their elevated C to T transition rate. Importantly, the protocol requires only a minimal departure from standard DNA-seq and, while the deamination is large enough to slightly elevate the C to T error rate at m5C sites, this level of deamination does not affect library yields and sequencing quality (Baum et al. 2021).

In this study, we developed RIMS-seq2 for the identification of methylated loci in humans. We first adapted the experimental protocol to assess concerted methylation at nearby CpG sites and apply RIMS-seq2 on well known cell lines as well as matched tumor/normal tissue samples. We demonstrate broad applicability of this technology in the simultaneous identification of sequence and methylation in a single experiment with minimal modification of the standard library protocol and sequencing qualities matching DNA-seq for variant calling.

## Result

### 1. Considerations for the application of RIMS-seq2 to human genome sequencing

A previous version of the RIMS-seq protocol has been used for the identification of methylase specificity in bacterial genomes using an overall elevation of C to T transitions rate of ∼ 0.1% (Baum et al. 2021). This deamination level is generally enough for the identification of sequence context(s) surrounding the methylated cytosine characteristic of the prokaryotic methylase specificity(ies). To apply RIMS-seq2 for the identification of human methylation, the protocol needs to be adapted to the fact that methylation happens almost exclusively at CpG sites and only a subset of these sites is methylated. Thus, the experimental and analytical pipelines need to be adjusted to identify regional methylation levels on the human genome.

Accordingly, we increase the signal by selecting conditions to achieve a ∼ 1% C to T transition at m5C sites. We also modified the previously published protocol to further lower the background of C to T transition at unmethylated sites **(Material and Methods)**. Under these conditions, the C to T transition is expected to increase to about 1000-fold at m5C sites compared to the background error rate of Illumina. This fold increase represents the goldilocks zone to identify methylation in human without affecting sequencing quality for a variety of standard genomic applications. For example, we expect the identification of germline variations to be done using standard tools for variant calling such as GATK (McKenna et al. 2010)) without affecting call accuracy.

Nevertheless, at current standard sequencing depth (30-fold coverage or above), this level of deamination does not provide base resolution methylation identification but rather provides regional aggregated methylation levels (RAML) over a defined genomic region. We estimated that the accurate evaluation of the methylation status can be safely done at about 100 combined CpG sites or above. Indeed, with 1% deamination rate and 30 fold coverage of a hundred CpG sites should theoretically yield a combined 30,000 reads and an average of 300 C to T transition events at methylated sites. This number of CpG sites represent about or less than the average size of a single CpG island, a resolution that is compatible with most epigenetics applications. Indeed, functional genomic regions ranging between a few hundred to a few thousand bases tend to be regionally methylated or unmethylated in concert (Chen, Lin, and Fann 2016) and a number of established protocols for methylation analysis have already been taking advantage of this concerted signal to identify such local aggregate as opposed to base resolution methylation levels (Weber et al. 2005)(Brinkman et al. 2010).

### 2. Whole Genome and targeted genome sequencing

To demonstrate the applicability of RIMS-seq2 in the simultaneous sequencing of DNA and methylome, we performed RIMS-seq2 on human genomic DNA. RIMS-seq2 whole genome was performed on NA12878 genomic DNA. Exome target capture using RIMS-seq2 were performed on both NA12878 and K562 genomic DNA as well as genomic DNA extracted from beast frozen tissue. Finally, to ensure reproducibility and compatibility with DNA-seq, we performed technical replicates on the target captures and included a control DNA-seq using the same source of starting material respectively.

For Whole genome sequencing, about 150 ng of NA12878 genomic DNA was used to generate 4.6 billion paired end reads, achieving an average coverage above 200x. For exome sequencing, we used 50-100 ng of NA12878 genomic DNA to generate about 100-200 million paired end reads, achieving an average of above 40-fold coverage **(Supplementary Table 1)**.

Reads were trimmed and mapped to the human genome (GRCh38) using bowtie2 (Langmead and Salzberg 2012). C to T transitions at CpG sites were identified for each individual reads and combined over pre-define genomic regions to obtain C to T transition rates (Material and Methods, Supplementary Table 1). Transition rates are subsequently calibrated to obtain overall methylation levels in these genomic regions (see below).

#### 2.1 RIMS-seq2 shows linear relationship between Transition rates and methylation level

We first evaluated how the C to T transition rate of RIMS-seq2 correlates with local methylation level in CpG Island (CGI), promoters or exonic regions. For this, we define methylation levels across the NA12878 genome using published gold standard datasets. More specifically, we calculated the weighted average methylation from three datasets derived from whole genome bisulfite sequencing (WGBS), EMseq (Vaisvila et al. 2021) and Nanopore (Jain et al. 2018) done on NA12878 **(Material and Methods)**. Because these datasets result from independent technologies for methylation identification, the weighted average methylation should minimize the bias inherent to each method (Olova et al. 2018) and provide a closer to the ‘true’ methylation levels (See methods). Next, CpG Island (CGI), promoters or exonic regions with similar methylation levels were binned together and the excess of C to T transition rate observed in RIMS-seq2 is computed for each bin.

As expected, we found the excess of C to T transition to be correlated with methylation levels in CGI **(Figure 1A)**, promoter and exonic regions **(Supplementary Figure 1A, and B)**. Such correlation was only observed in CpG context, the other contexts do not show excess of C to T transition consistent with the fact that the vast majority of methylation events in human happen in CpG context (Figure 1A).

**Figure 1:**
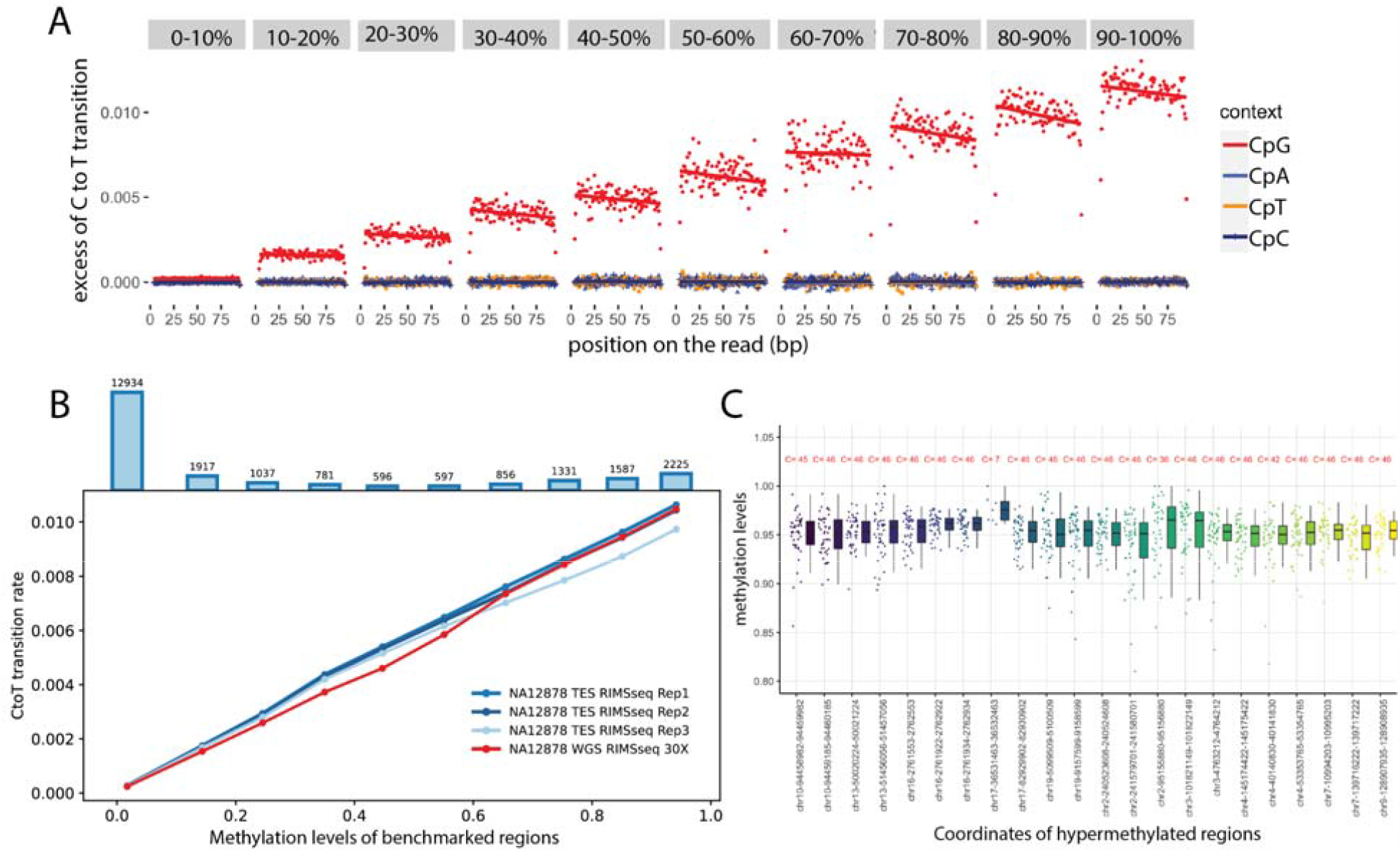
linear relationship between Transition rates and methylation level. **(A)** Whole genome RIMS-seq2 excess of C to T transition rates in read 1 compared to read 2 (Imbalance) function of the position on the read (in bp). CGI regions are binned into 1-10% to 90-100% methylation levels. The rate of C to T transition in each bin was computed for CpG (red), CpA (blue), CpT (orange) and CpC (brown) contexts. **(B)** The rate of C to T transition and benchmarked methylation in binned CGI fit a positive linear regression model for all NA12878 RIMS-seq2 whole genome (red) and exome sequencing in triplicates (blue). Top bar plots represent the total number of CGI in each bin **(C)** Twenty-two stably methylated regions across a broad range of tissues and cell types used as internal control for methylation level calibration. The number in red indicates the number of available human WGBS samples used for methylation analysis. Genomic regions coordinates are derived from the human genome (Assembly GRCh38).

To demonstrate linearity between methylation levels and RIMS-seq2 C to T transition rates, we performed a regression analysis test for the binned CGI **(Figure 1B-C, Supplementary Table 3)**, promoters and exons **(Supplementary Figure 1C and D)** for both whole genome and targeted RIMS-seq. The resulting regression analysis reveals positive linear relationships between transition rates and the methylation levels in all regions for both whole genome (R square = 0.99, p value of variable < 0.01, Coefficient = 0.01) and targeted RIMS-seq2 (R square = 0.99, p value of variable < 0.01, Coefficient = 0.01). This result indicates that quantitative measurement of methylation level using RIMS-seq2 is achievable and the absolute quantification can be calibrated using only two data points.

#### 2.2 Development of a linear model between transition rates and methylation levels

While the heat alkaline deamination treatment is aiming at about 1% C to T transition at m5C sites, experimental variations, genomic DNA quality and sequencing runs may affect the C to T transition rate at both C and m5C for each individual experiment. It is therefore important to perform a calibration specific to each sequencing run. Having established the linear correlation between transition rate and methylation levels, this calibration can be done using two sets of internal controls only representing hyper methylated regions and unmethylated sites respectively.

For hyper methylated regions, a set of 24 internal control regions were selected as stably hypermethylated across a broad range of tissues and cell types (Edgar et al. 2014) including in NA12878 **(see Material and Methods and Figure 1C)**. We used the non-CpG sites in these regions to estimate the C to T transition rate at unmethylated cytosines. Applied to all the RIMS-seq2 datasets obtained for this study **(Supplementary Table 1)**, we obtained C to T transition rate at m5C to be around 1% for m5C and 10.e-5% for C, that is a 1000-fold increase in C to T transition rate at methylated compared to unmethylated sites.

Next, using the stably hypermethylated regions, we investigated whether sequence context matters for calibration, notably the upstream base relative to the deaminated base. The effect of the upstream base to CpG sites is subtle but not negligible with sample and run variations **(Supplementary Figure 2A)**. ApCpG context for which an adenosine is found upstream of the CpG repeatedly shows the lowest C to T transition **(Supplementary Figure 2A)**. The sequence context is therefore incorporated into the calibration procedure as a parameter to improve methylation call accuracy. We also observed an effect of the sequencing cycles, notably for the two first and two last cycles of paired-end sequencing **(Supplementary Figure 2B)**. For these cycles, the C to T transition is lower than expected for fully methylated sites and removed from the methylation analysis.

Finally, since the deamination rate only moderately increases the signal over the noise ratio, RIMS-seq2 is attuned to variations from the reference genome. Thus, we also investigated the effect of single nucleotide polymorphism (SNP) on methylation calls. We observed an increase in methylation call accuracy if the publicly available NA12878 SNPs positions were to be removed prior to methylation call **(Supplementary Figure 2C)**. An equivalent improvement was obtained if the SNPs positions were identified directly from the RIMS-seq2 datasets and subsequently used for methylation call (see below for details) **(Supplementary Figure 2C)**. This result demonstrates that an external SNP dataset is not required for this analysis. Thus, prior to calibration, the positions identified by RIMS-seq2 as SNPs were removed from the methylation calls.

### 3 Methylation calling at regional resolution

We are now addressing the ability of RIMS-seq2 to define methylation at regional resolution. As a first pass, profiles of C to T rate were compared to a published WGBS methylation profile performed on GM12878. Visual inspection of the methylation profile indicates that the C to T profile correlates closely with the methylation level profiles from BS sequencing **(Figure 2A)**.

**Figure 2:**
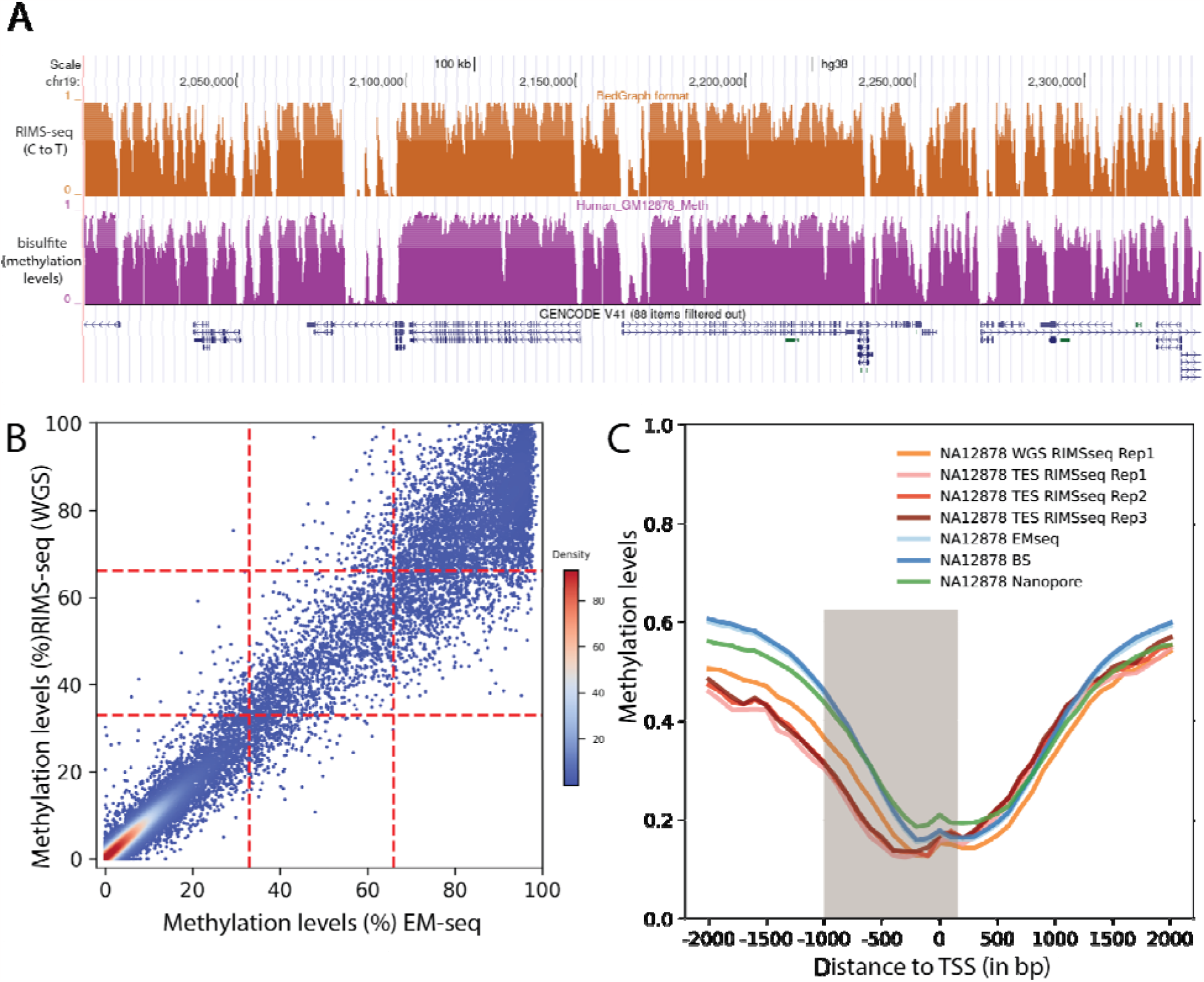
Performance of RIMS-seq2 (Methylation). **A**. C to T profile at a specific locus (combined 30 CpG sites) compared to methylation profile from Bisulfite sequencing **B**. <u>Methylation level (RAML) correlation at CGIs between RIMS-seq2 and EM-seq</u>. Each point corresponds to a CGI region. The plotting area has been divided into 9 quadrants to reflect the correlation. **C**. <u>Methylation profiles at promoters</u> measured by RIMS-seq2, EM-seq, Bisulfite sequencing (BS) and Nanopore. The overall percentage methylation of CpG sites were measured using 100 bp sliding windows within 2kb upstream and downstream of transcription start sites (TSS). TSS were defined using USCS annotation. Distance to TSS is measured in bp. The shaded region corresponds to the definition of promoters (-1kb downstream and 100 bp upstream of TSS respectively)

To quantify how correlated the methylation levels are between RIMS-seq2 and other technologies such as bisulfite sequencing, EM-seq and Nanopore, we proceed with a genome-wide comparison of various publicly available methylation datasets done on the same cell lines. Since RIMS-seq2 cannot reliably identify methylation at base resolution with current standard sequencing depth, we aim at obtaining regional aggregated methylation levels (RAML) values at defined genomic regions. For this, we delineated genomic regions of interest for methylation identification such as CpG island (CGI), promoters and exonic regions and performed local calibrations using the linear model described above for the combined CpG sites in these regions. For comparison, we also perform similar regional methylation aggregates with the public methylation datasets. We found that the large majority of CGIs (67%) and promoters (70%) have methylation levels below 30% **(Figure 2C)** indicating hypomethylation in these regions consistent with the fact that these regions tends to be hypomethylated (Weber et al. 2007). Conversely, only 30% of exonic regions are hypomethylated.

For comparison with existing technologies, we performed standard correlation coefficients and complemented the correlation coefficients with a measure of quadrant consistency for an additional metric of similarity. Technical replicate analysis of RIMS-seq2 performed on the same sample shows good methylation concordance with quadrant consistency of 91% and a 0.95 correlation **(Supplementary Figure 3)**. The same correlation levels are observed between triplicate RIMS-seq2 exome sequencing and WGS **(Supplementary Figure 3)**. High correlation levels are observed in replicates of both cell lines as well as the frozen tissue **(Supplementary Figure 3, Supplementary Figure 4)**. Importantly, the correlations between RIMS-seq2 and other technologies are similarly high ranging from 0.94 to 0.98 **(Figure2B, Supplementary Figure 4)**.

### 3. Comparison with available technologies for genome sequencing

#### a. Coverage bias, insert sizes, chimeras and on-target sequencing

We perform basic quality control on both the RIMS-seq2 dataset and standard DNA sequencing. **(Supplementary Figure 5)**.

#### b. Germline variant calling

The deamination conditions should not interfere with genome sequencing for a variety of applications such as the identification of germline variations. To demonstrate that RIMS-seq2 accurately identifies germline variation, we use the GATK pipeline (McKenna et al. 2010) for variant calling on both the whole genome and exome RIMS-seq2 data and compare the results to JIMB variants focusing on SNPs. If deamination is interfering with variant calling, the overall fraction of C to T or G to A transition should be higher in RIMS-seq2 datasets. Nonetheless, the profile of SNPs closely resembles the JIMB SNPs indicating that the overall SNPs profiles are not affected **(Figure 3A)**.

**Figure 3:**
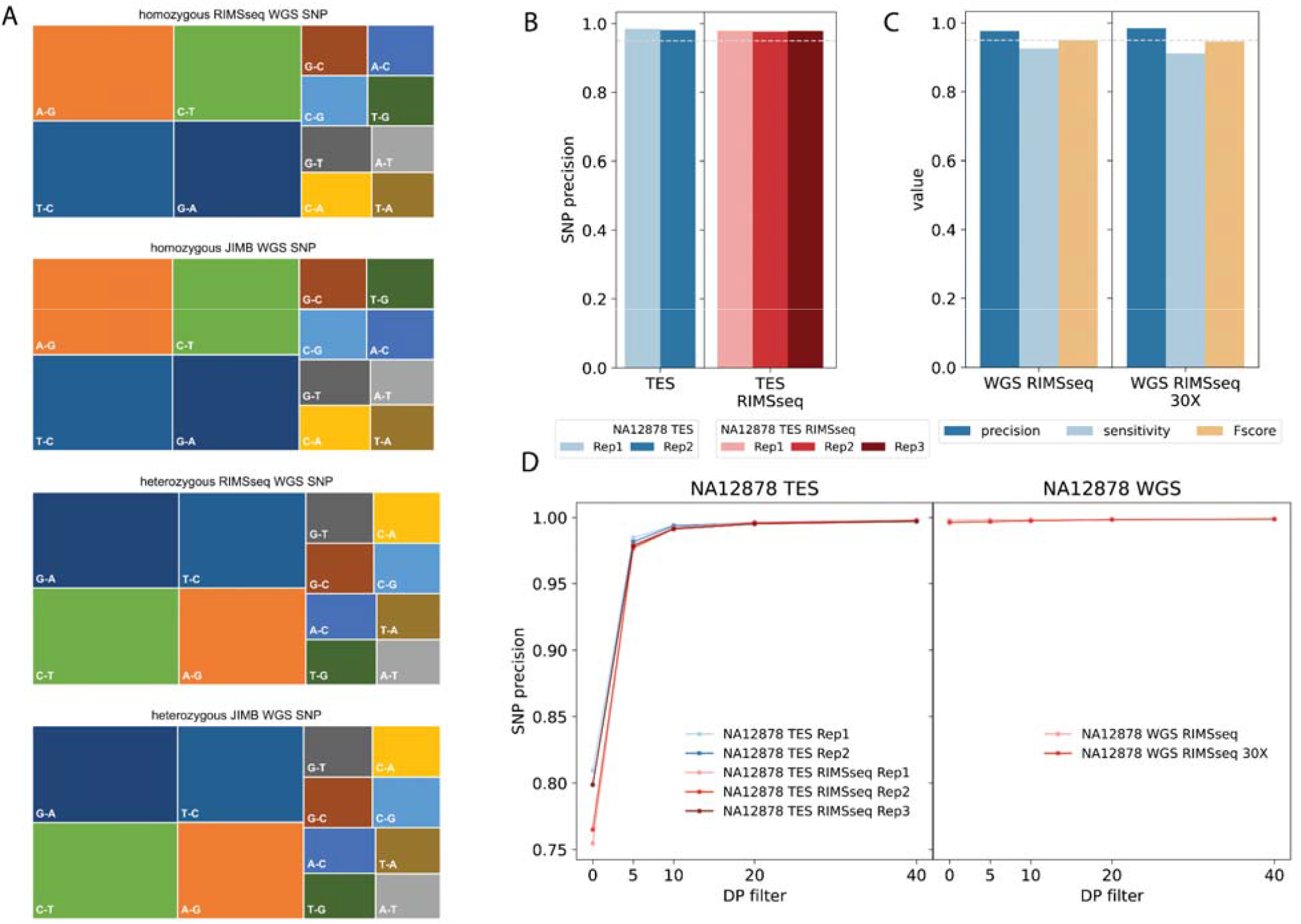
Performance of RIMS-seq2 (DNA sequencing). **A**. Germline transitions/transversion profiles for JIMB (left) and RIMS-seq2 (Right) WGS for homozygous (top) and heterozygous (bottom) SNPs. In all cases, transitions represent the majority of the SNPs identified. **B**. Precision of variant calling for standard DNA-seq and RIMS-seq2 targeted exome sequencing. **C**. Precision, sensitivity and F-score for WGS RIMS-seq2 (full dataset and downsampled to 30 fold coverage). **D**. ROC curve.

Finally, we apply RIMS-seq2 to the detection of methylated regions in frozen tissue samples. Exome sequencing was performed in triplicates and compared to EM-seq performed in duplicate on the same sample (Material and Methods). A correlation of an average of 0.95 can be observed between RIMS-seq2 and EM-seq **(Supplementary Figure 4)** demonstrating that RIMS-seq2 can also be used on more complex samples such as frozen tissue samples.

## Discussion

RIMS-seq2 enables the simultaneous identification of sequence and methylation for short read sequencing. Importantly, the experimental setup closely resembles a standard library preparation with minimal changes and the resulting data can be analyzed using standard variant calling. These features make this technology an extremely easy to deploy strategy for simultaneous germline variation and methylation identification in large scale sequencing labs.

Simultaneous germline variation and methylation identification is now routinely performed using long-read sequencing (Rand et al. 2017)(Simpson et al. 2017) but only in the context of native DNA for which no prior amplification has been performed, limiting the range of applications for direct methylation sequencing. Conversely, RIMS-seq2 can be performed with or without amplification and thus, can be performed on a greater set of applications.

At current standard sequencing depth of 30-fold for whole genome sequencing, RIMS-seq2 cannot be base resolution. Nonetheless, with constant increase in throughput and price drops, it is conceivable that several thousand-fold coverage of the human genome can be achievable in a routine fashion. Such coverage levels are already possible for targeted genome sequencing enabling base resolution methylation calls using RIMS-seq2.

Importantly, RIMS-seq2 is compatible with a large number of pre-sequencing treatments such as target enrichment as demonstrated in this study but also chromatin accessibility sequencing, ChIP-seq or single cell. In conjunction with target enrichment, quality control metrics show essentially identical performance compared to the DNA-seq.

As we have demonstrated in this study, the 1% deamination at CpG sites does not interfere with germline variation calls because variant frequencies are significantly above the deamination rates. Nonetheless, the same cannot be said for the identification of rare somatic mutations for which the frequency of variant is similar to the deamination rate. In these cases, deamination may confound the identification of rare somatic mutations and may not be used for these applications. Alternatively, algorithms for somatic mutations can be adapted to distinguish true mutations from limited deamination.

We have shown that the sequencing context has a minimal effect on the deamination rate of methylated cytosine. Thus, RIMS-seq2 can be directly applied to the identification of methylation in organisms that methylate cytosines in other contexts than CpG context such as plant and prokaryotes. Likewise, in this work, we choose to focus our analysis on CpG sites but CpA sites can be included for samples that are known to have significant levels of CpA methylation.

## Acknowledgments

We thank the NGS core sequencing group and Peter Weigele and Yan-Jiun Lee for providing purified XP12 phage genomic DNA.

## Data and code availability

All raw and processed sequencing data generated in this study have been submitted to the NCBI Gene Expression Omnibus (GEO; https://www.ncbi.nlm.nih.gov/geo/) under accession number GSE234235.

## Supplementary Table legends

Supplementary Table 1 : Summary of the DNA-seq and RIMS-seq2 sequencing (number of paired-end reads, mapping efficiency, PCR duplicates and other QC metrics from picard CollectHsMetrics.

Supplementary Table 2 : Regression analysis.

Supplementary Table 3 : 24 ultra-stably methylated regions in the human genome (GRCh38 coordinates).

## Material and Methods

### RIMSseq library preparation

We used human genomic DNA isolated from GM12878 cells (NA12878, provided by Coriell Institute), K562 cells (provided by ATCC) and breast tissue (Biochain D8235086-PP-10) for RIMSseq sequencing in this study. We used 100-200 ng and 50-100 ng of genomic DNA for RIMSseq whole genome sequencing (WGS) and exome targeted sequencing (TES) library preparation, respectively. We performed RIMSseq following the published protocol (Baum et al. 2021) with some modifications. We used NEBNext Ultra II library prep kit (NEB E7645) following the manufacturer’s instructions until the USER treatment step included following the adapter ligation step. After this first USER treatment step, the sample was subjected to heat alkaline deamination in 1 M NaOH (final concentration) for 30 min at 60⍰.The sample was subsequently cooled down on ice and an equal moles of acetic acid was added to a final concentration of 1 M to neutralize the PH. DNA was purified using Zymo oligo clean and concentrator kit (D4060 Zymo Research) following the protocol for clean-up of DNA above 80 nt. An additional USER treatment step was performed to the purified DNA by adding 2μl USER (included in all the NEB index primer kits) and incubating for 15 min at 37⍰. Finally, we used the USER treated DNA as template for PCR amplification using NEBNext Ultra II Q5 Master Mix. Eight samples were amplified and pooled for target enrichment using the Twist Comprehensive Exome Panel (Twist 102031), following the manufacturer’s recommendations. The enriched DNA was subsequently amplified with NEBNext Ultra II Q5 Master Mix, and both the whole genome and targeted libraries were sequenced on the Illumina Novaseq 6000 platform using a paired-end mode with a read length of 100 base pairs.

### EM-seq library preparation

Fifty nanograms of genomic DNA from breast tissue was used to prepare EM-seq libraries as per the manufacturer’s instructions (NEB E7120). The Illumina Novaseq 6000 sequencer was used to sequence the libraries in a paired-end mode, generating 100 base pair reads. The evaluation of EM-seq conversion efficiency (>99.7%) was performed by utilizing unmethylated lambda genomic DNA as a spike-in.

## Data analysis

### Reference genome and other annotation files

We used the GRCh38 human reference genome (hg38), UCSC human CpG island annotation and known human SNP files used for GATK Base Quality Recalibration as explained in (Yan et al. 2022).

### RIMSseq Data processing

Initially, Trimgalore (version 0.6.4) was utilized to trim the Illumina adapter from the reads. Additionally, for NA12878 WGS RIMSseq, the first two bases of Read1 were trimmed owing to their poor quality (--clip_R1 2). Next, the trimmed reads were aligned to the hg38 human reference genome using Bowtie 2 (version 2.3.0) with the default parameters for paired-end mapping and inclusion of the read group identifier defined by @RG. To ensure the accuracy of downstream analysis, we discarded improperly mapped reads using SAMtools (version 1.14) and PCR duplicates using Picard tools (version 2.26.11) MarkDuplicates.

### RIMSseq C to T transition counting

To prevent the repetitive counting of the same transition event, we utilized a custom script (Github: TrimOverlappingReadPair.py) to remove the overlapping regions between Read1 and Read2 from Read2. Next, we separated the mapped Read1 (-f 64) and Read2 (-F 64) using SAMtools. Then we compared the Read1 and Read2 mapping to the hg38 genome using SAMtools mpileup with the following parameters: --min-MQ 10 --min-BQ 30 --output-BP-5 --no-output-ins --no-output-ins --no-output-del --no-output-del --no-output-ends. The C to T transition at CpG sites was then counted for both Read1 and Read2 in a context-dependent manner using a custom script CountErrorMpileup.py (Github), with the following parameters: -- REF C --BASE T --left 1 --right 0 for Read1 and --REF G --BASE A --left 0 --right 1 for Read2. Context-independent counting was performed using --left 0 --right 0 for both Read1 and Read2. Furthermore, the removal of SNP positions and specific sequencing cycles from counting was accomplished using the --vcf and --cycle options, respectively, to enhance the accuracy of downstream methylation prediction, as discussed in the figure XX. Finally, we added the C to T transition of all the CpG sites in the targeted region(s) such as CGI using a custom script CountErrorRegion.py (Github). These regional C to T transition counts (defined as Error) and all cytosine counts (defined as Total) were used for regression analysis and methylation prediction. The C to T transition rate (R) equals to Error divided by Total.

### Data processing and methylation quantification by other methods

We downloaded the previous published EMseq or WGBS data set: NA12878 ENCODE WGBS (ENCODE, ENCSR890UQO), NA12878 NEB WGBS (NCBI SRA, SRR10532136, SRR10532135, SRR10532127 and SRR10532126)(Vaisvila et al. 2021), NA12878 NEB EMseq (NCBI SRA, SRR10532145, SRR10532144, SRR10532139 and SRR10532138) (Vaisvila et al. 2021) and K562 ENCODE WGBS (ENCODE, ENCSR765JPC). The EMseq of breast tissue was generated in this study as mentioned above.

We processed these data sets and extracted methylation information using the bismark pipeline: (1) for a fair comparison, we shortened the paired-end reads to 100 bp long and trimmed the Illumina adapters as well as the first two bases of Read2 (Trimgalore --clip_R2 2); (2) we aligned trimmed reads to the human GRCh38 genome using Bismark (version 0.22.3); (3) we filtered the PCR duplicates and incomplete bisulfite conversion using Bismark deduplicate_bismark and filter_non_conversion, respectively; (4) we combined the replicates and extracted CpG sites methylation using bismark_methylation_extractor. The methylation level of targeted regions such as CGI or promoters was calculated as explained previously (Yan et al. 2022). For NA12878 BS-tagging sequencing (mentioned as Tn5 BS in Figure) (Yan et al. 2022; Suzuki et al. 2018), we converted its processed methylation information on hg19 to hg38 using the UCSC liftOver tool (Hinrichs et al. 2006). For NA12878 Nanopore sequencing, the data processing and methylation quantification were described in (Yan et al. 2022).

### Regression analysis between the C to T transition rate and the methylation level

We mathematically described the relationship between the RIMSseq C to T transition rate and the methylation level of CpG sites in certain regions of the GM12878 cells. We chose the CpG islands, promoter regions that are defined as 1000 bp upstream and 100 bp downstream of the annotated TSS and Twist target enrichment exome bait regions for this analysis. The methylation level of these regions was measured by three methods including whole genome bisulfite sequencing, EMseq and Nanopore sequencing as mentioned above. We used regions having coverage of CpG sites >=50 in WGBS and EMseq and >=20 in Nanopore for analysis.

To reduce method bias, we integrated the measurements from these three methods, which is defined as the benchmarked methylation level for the region n (BMn), using the following steps: (1) we calculated the proportion (*P*_*n,i*_) and weight (*w*_*n,i*_) of each method for region n given by

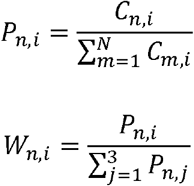

Where *i, j∈* {1,2,3} denote the three methods; *n,m* {1,2, …,*N*} denote the region and C _*n,I*_ represents the coverage of CpG sites by the method in certain region n; (2) For region n, we selected the two measurements with closest values given by

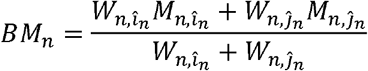

Where *M*_*n,i*_ and *M*_*n,j*_ stand for the methylation level measured by method *i* and *j* for region n. We computed the BMn which is the weighted average methylation by

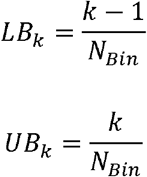

Next, we classified these regions into *N*_*Bin*_ bins (*N*_*Bin*_ = 10) with equal width based on the benchmarked methylation level. The lower bound (*LB*_*k*_) and upper bound (*UB*_*k*_) of each bin is defined as

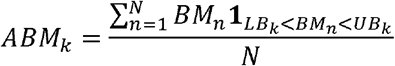

with *k ∈* {1,2, …,10}. The average methylation level of bin *k* (ABM_*k*_) is calculated by

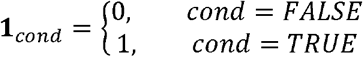

where N is the number of CGI regions in bin *k* and represents the indicator function. For a given condition,

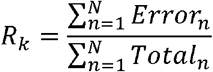

therefore *BM*_*n*_1 *LB*_*k*_ <*BM*_*n*_<*UB*_*k*_ means the benchmarked methylation within the lower and upper bound. We also measured the RIMSseq C to T transition rate with the following parameters as explained above: --min-MQ 10 for samtools mipleup; --vcf using SNP annotation based on JIMB whole genome sequencing of NA12878 for counting error using CountErrorMpileup.py. Then the total cytosine counts (*Total*_*n*_) and C to T transition counts (*Error*_*n*_) in all the regions included in bin *k* were added together to represent the transition rate (*R*_*k*_) of this bin

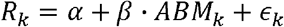

with N is the number of regions in bin *k*.

Finally, we performed the regression analysis to evaluate the linear relationship between the methylation level *ABM*_*k*_ and transition rate *R*_*k*_. By interpreting the P-value of variable and intercept, we concluded that these two variables fit the linear model as shown in Figure X and Supplemental Table X

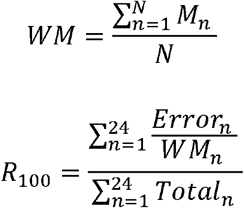

### RIMSseq regional aggregated methylation level (RAML) quantification

Given the methylation level and C to T transition rate fitting a linear model, we can predict the regional aggregated methylation level based on the C to T transition rate of the target region. We used 24 hypermethylated regions (Supplemental Table X), for which the CpG sites are known and confirmed to be stably methylated in the human genome, to establish the linear model. We use *R*_0_ and *R*_100_ to represent the C to T transition rate of the non CpG cytosines and the CpG sites in these stably hypermethylated regions, respectively. It is worth mentioning that we adjusted *R*_100_ corresponding to the real methylation level in these regions based on all the published human WGBS data from ENCODE as annotated in Supplemental file, given by

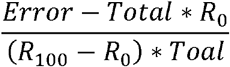

Where *M*_*n*_ represents the methylation level of a certain hypermethylated region based on one human

ENCODE WGBS dataset, and N is the number of available human WGBS datasets for this region. Therefore, *WM* represents the mean methylation level of this hypermethylated region in human genome.

The RAML can be estimated by

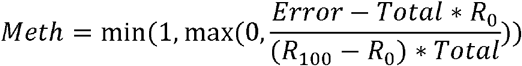

*Total* and *Error* are the cytosine counts and C to T transition counts in the corresponding region as explained above. Since the methylation level needs to be between 0% and 100%, we apply a lower and upper bound for the estimated methylation level (*Meth*)

[uneq]

### Variant calling and SNP comparison

The variant calling and comparison were performed as described previously (Zhou et al. 2019; Yan et al. 2022), except for the use of GATK version 4.2.5.0. For RIMSseq WGS and TES, we applied an additional filter “DP < 5” to remove SNPs with low coverage. The resulting SNP sites were used in the RIMSseq methylation prediction process. We used the WGS of NA12878 (generated by JIMB NIST Genome in a Bottle, JIMB WGS HG001 and K562 (ENCODE, ENCSR053AXS)(Zhou et al. 2019) as benchmark for variant calling comparison. We restricted the comparison to the variants on somatic chromosomes and Chr X.

---

**Supplementary Figure S1:**
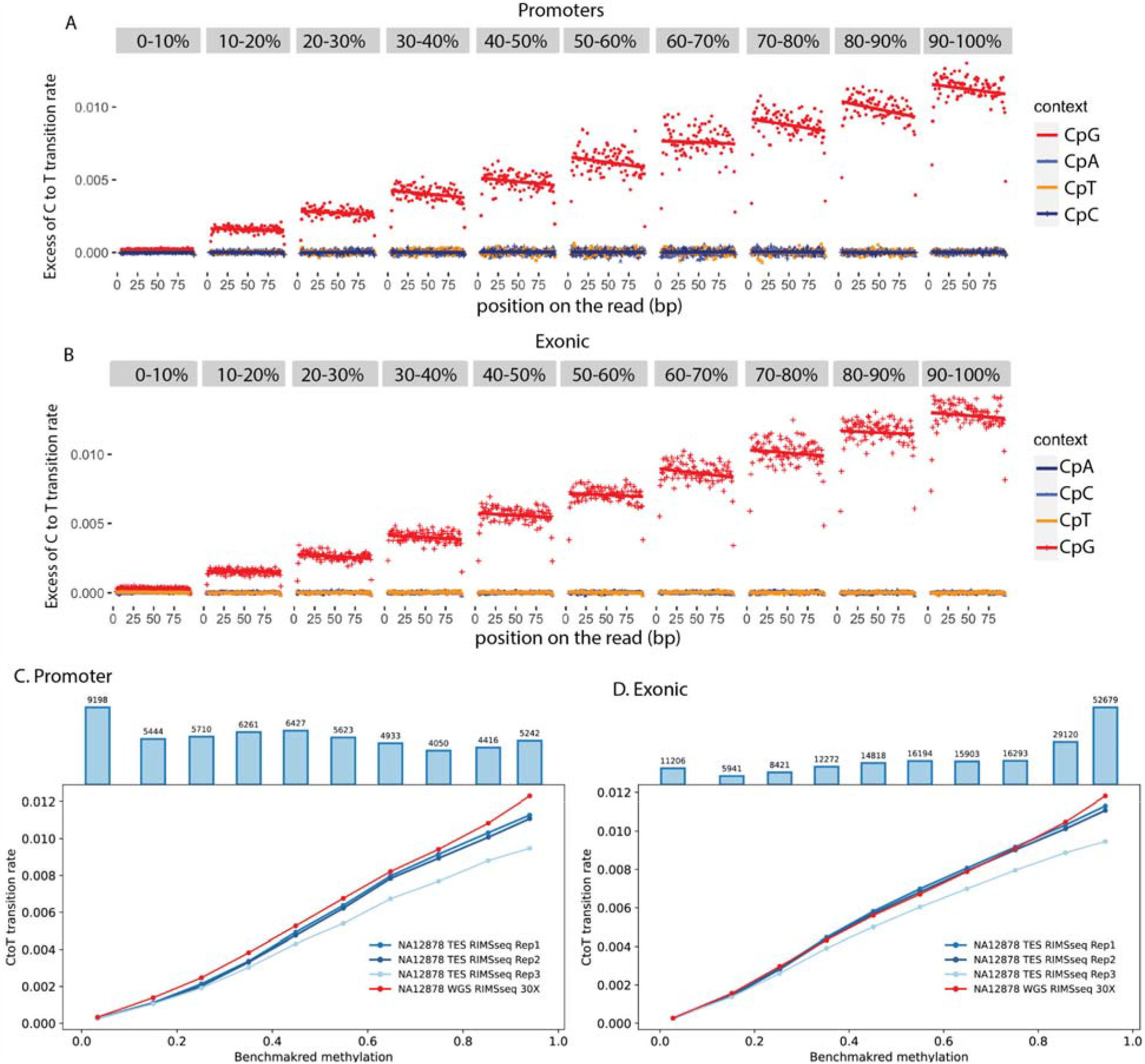
Whole genome RIMS-seq2 excess of C to T transition rates for **(A)** Promoter regions and **(B)** exonic regions binned into 1-10% to 9-100% methylation levels function of the position on the read. The rate of C to T transition was computed for CpG, CpA, CpT and CpC contexts. The rate of C to T transition and benchmarked methylation fit a positive linear regression model for **(C)** Promoter regions and **(D)** exonic regions in Whole genome RIMS-seq2 (red line) and all targeted RIMS-seq2 (blue lines). Blue bar plots represent the number of genomic regions for each binned methylation level.

**Supplementary Figure 2:**
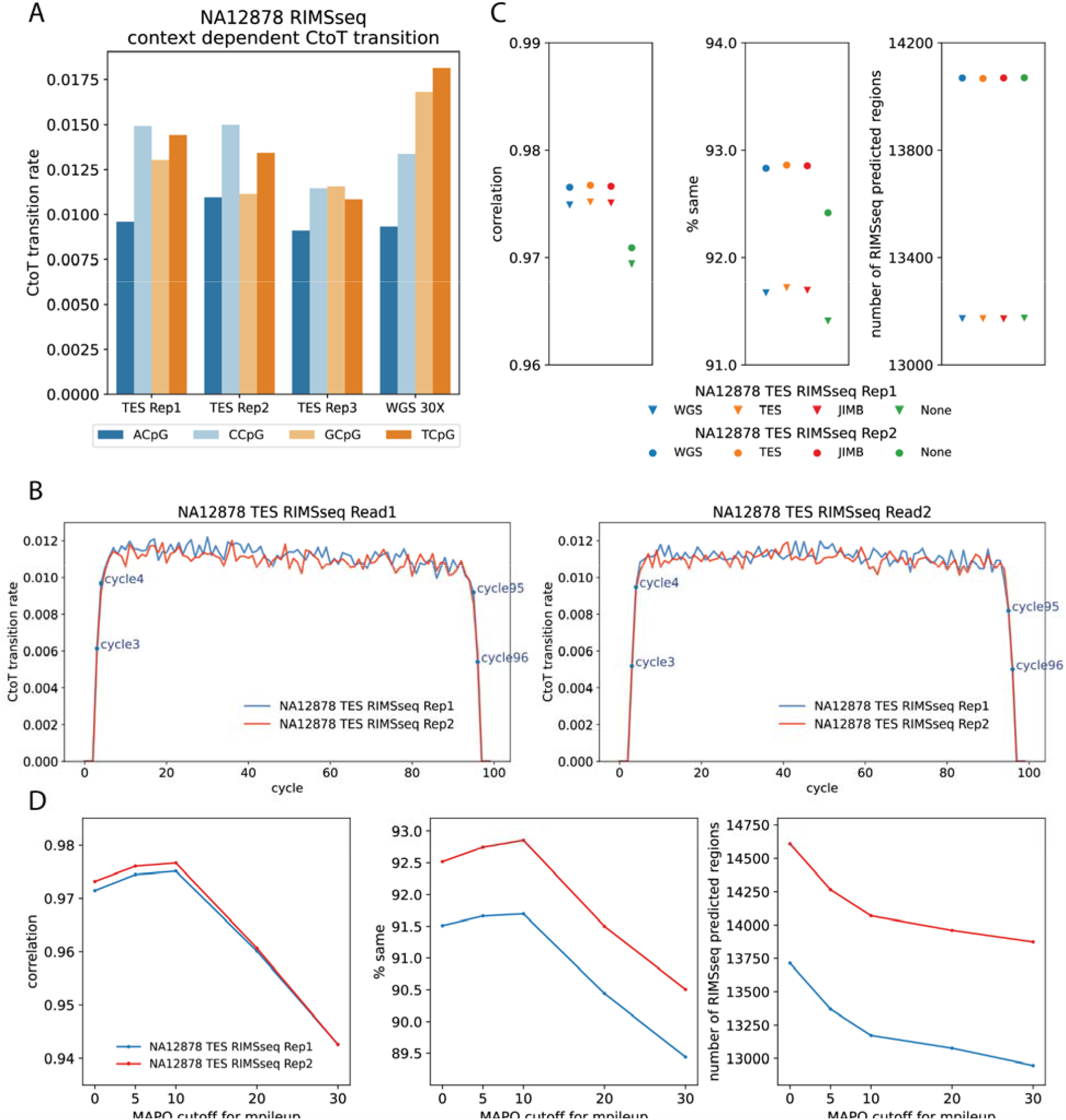
Additional parameters affecting calibration. **A**. <u>Sequence context</u> : C to T transition rates in the stably hypermethylated regions for the ApCpG (blue), CpCpG (light blue), GpCpG (light orange) and TpCpG (orange) sequence context for the three RIMS-seq2 exome sequencing replicates (TES Rep1, 2 and 3) and whole genome sequencing downsampled to 30 fold coverage (WGS 30x) **B**. <u>sequencing cycle</u> : C to T transition rates in the stably hypermethylated regions function of the sequencing cycles for two exome replicates for the first (left) and second (right) paired end read. **C**. <u>SNP</u> : Confounding effect of germline variation on methylation calls **D**. <u>MAPQ cutoff</u> : effect of MAPQ cutoff on methylation calls.

**Supplementary Figure 3:**
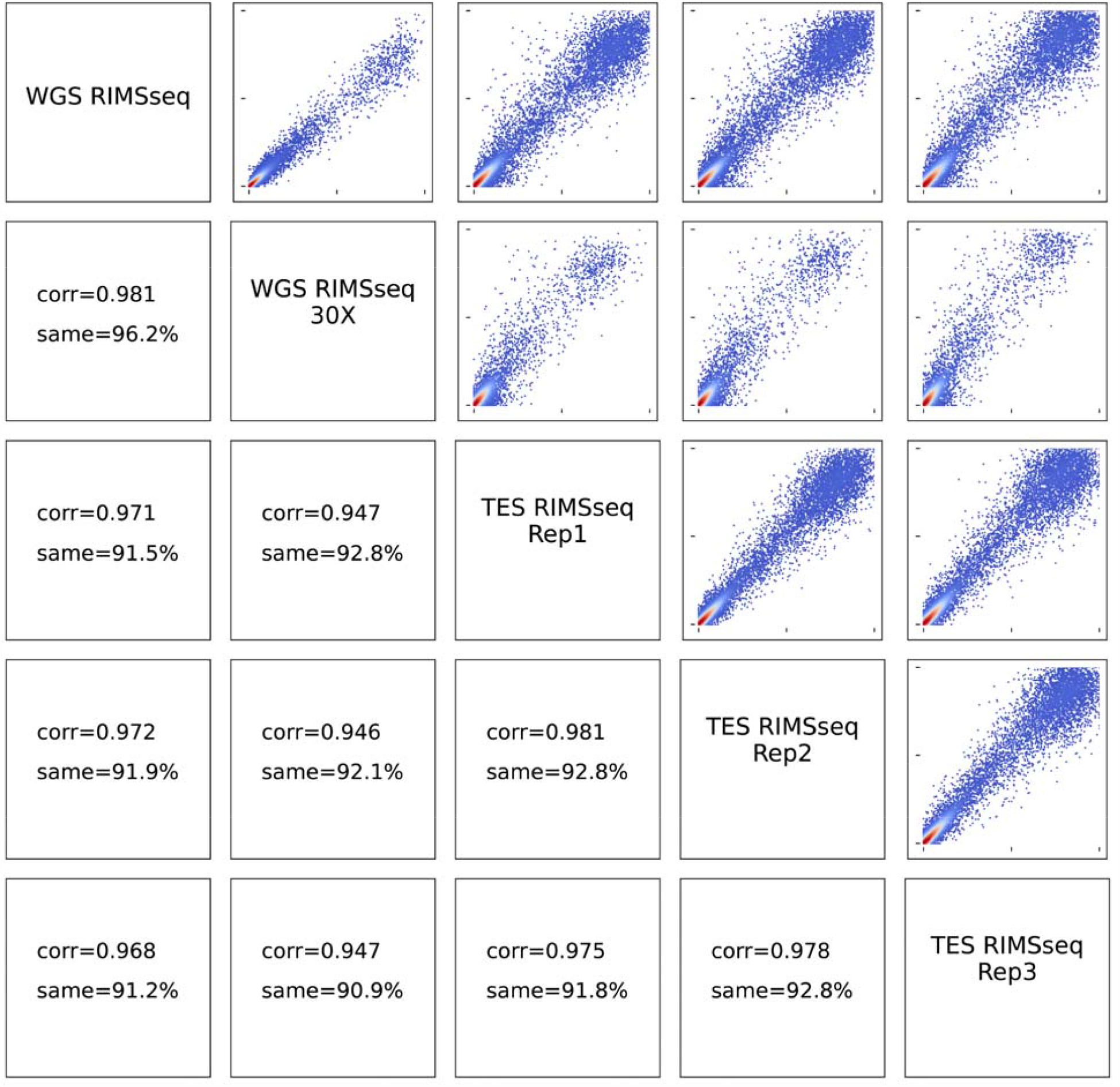
RIMS-seq2 replicate correlations. Comparison of the methylation levels of CGI (RAML) for one whole genome sequencing (WGS, full dataset and downsampled to 30x coverage) and exome sequencing performed in triplicates (TES Rep1, Rep2 and Rep3). Corr stands for Pearson correlation. Same represents the percent of CGI that are quantified in the same methylation category (high, medium or low) by both methods.

**Supplementary Figure 4:**
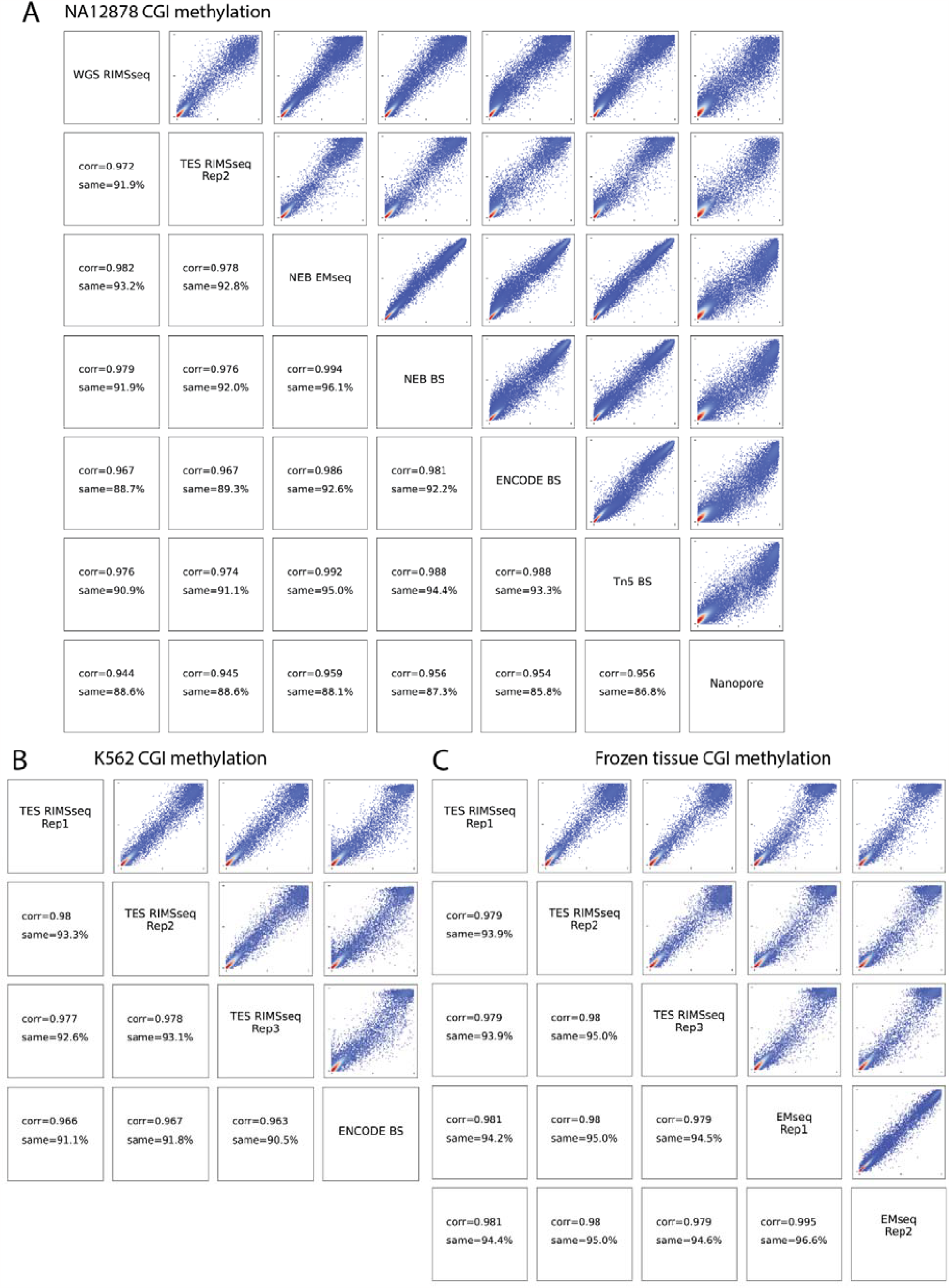
RIMS-seq2 correlations with other sequencing technologies. CGI methylation quantification of **A** NA12878 **B**. K562 **C**. Frozen tissue. BS: bisulfite. Tn5 BS: Bisulfite-tagging sequencing. Corr stands for Pearson correlation. Same represents the percent of CGI that are quantified in the same methylation category (high, medium or low) by both methods.

**Supplementary figure 5:**
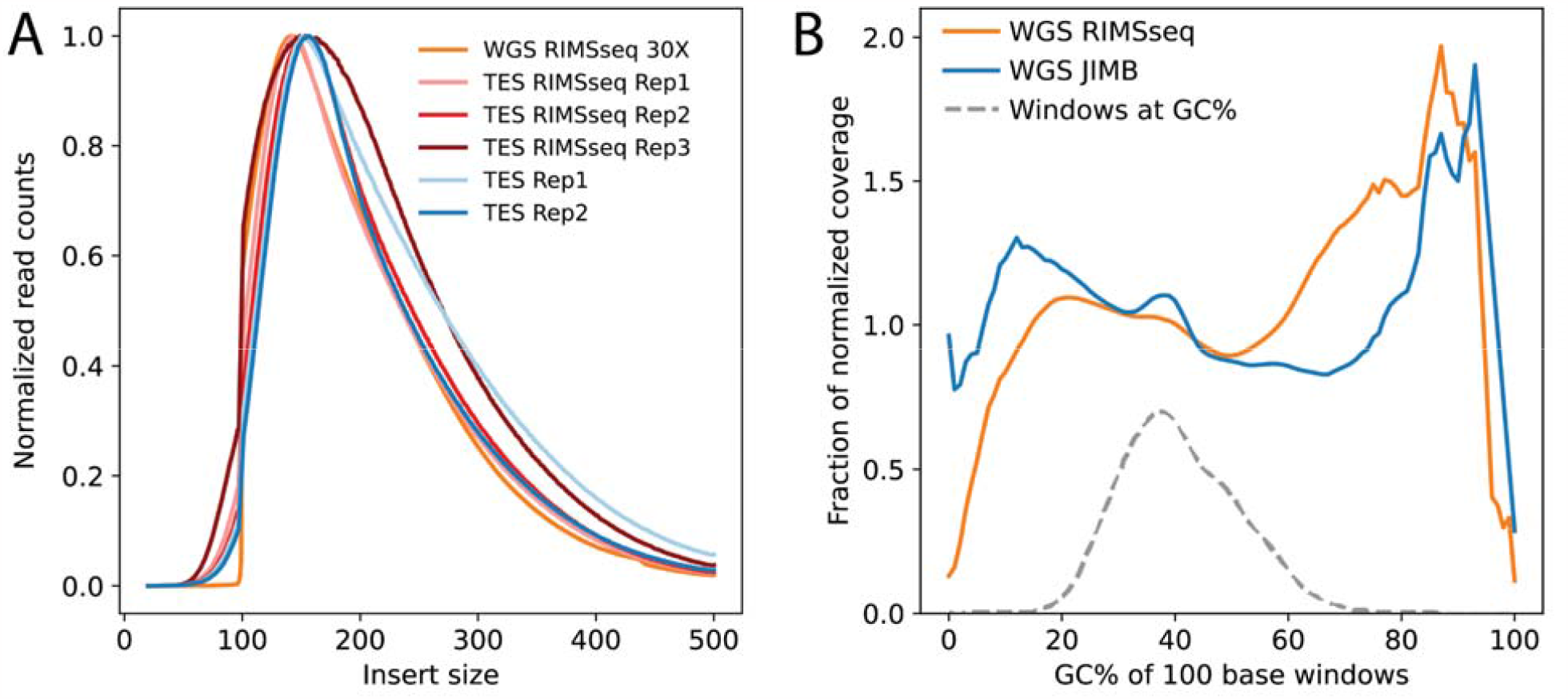
Standard quality control for DNA-seq. **A**. Insert size distribution (in bp) for RIMS-seq2 libraries (red, orange, pink) and DNA sequencing (blue) **B**. CG bias for the RIMS-seq2 (orange) and DNA-seq (blue) whole genome sequencing.

